# HerpesDRG: a comprehensive resource for human herpesvirus antiviral drug resistance genotyping

**DOI:** 10.1101/2020.05.15.097907

**Authors:** OJ Charles, C Venturini, RA Goldstein, J Breuer

## Abstract

The prevention and treatment of many herpesvirus associated diseases is based on the utilization of antiviral therapies, however therapeutic success is limited by the development of drug resistance. A comprehensive point of truth of resistance conferring mutations has been missing but would be important to aid the development of antiviral drugs and in management of infections. We therefore developed HerpesDRG, a drug resistance mutation database for all the known important genes and current treatment options, built from a systematic review of available genotype to phenotype literature. The database is released along with an R package to provide a low barrier of entry to variant resistance annotation and clinical implication analysis from common sanger and NGS sequence data. This represents the first openly available and community maintainable knowledgebase of drug resistance mutations for the human herpes viruses (HHV), developed for the community of researchers and clinicians tackling HHV drug resistance.

**Availability:** The HerpesDRG database is available at github.com/ojcharles/herpesdrg-db. The R package for resistance genotyping data is available at github.com/ojcharles/herpesdrg. A user-friendly webserver is available at cmv-resistance.ucl.ac.uk/herpesdrg

## Introduction

Herpesviruses and their associated diseases are a major health problem worldwide for immunocompromised patients. The prevention and treatment of human cytomegalovirus (HCMV) and herpes simplex virus (HSV) disease for example are essential in management of solid organ transplant (SOT) & hematopoietic stem cell transplant (HSCT) recipients (Ariza-Heredia et al., 2014; Emery, 2001; Lee et al., 2019; Limaye, 2002). Although a handful of treatment options are available such as Acyclovir, Ganciclovir and Letermovir, emergence of resistance causing mutations are now found for all approved drugs and pose a significant threat due to aggressive disease course and a greater mortality risk (Bacon et al., 2003; Kakiuchi et al., 2017; Komatsu et al., 2014; Lurain and Chou, 2010; Patel et al., 2014).

Drug resistance in a clinical setting is often diagnosed by sequencing followed by variant annotation (genotyping), rather than isolating and phenotypically characterising (phenotyping) due to cost and time benefits (Gu et al., 2019; Russell, 2018). Genotyping does however require that any variants present have previously been phenotyped (Lurain and Chou, 2010). In these experiments a single novel mutation is transferred into a reference virus, or a clinical isolate is found with similar characteristics. The impact of this mutation on drug sensitivity can then be determined by a plaque reduction assay (PRA), Bacterial Artificial Chromosome (BAC) reporter assay or similar, returning an IC_50_ against a given drug. It is typical to report the mutations impact on antiviral sensitivity as a fold change relative to the control strain IC_50_, where HCMV is currently the only human herpes virus (HHV) with consensus guidelines relating fold change to a “resistant” or “susceptible” phenotype (Kotton et al., 2018; Danve et al., 2002; Chou et al., 2005; Sauerbrei et al., 2011).

An issue is that those previously phenotyped data are published in a disparate collection of reports. This has meant that researchers often must perform a review of the scientific literature for identified mutations in a viral isolate, which is time consuming, especially when there are several reports for a given mutation (Paolucci et al., 2021; van der Beek et al., 2013, p. 20). Therefore there have been published efforts to collate this data previously such as in reviews, however the data contained is not updateable and requires reformatting for informatics (Lurain and Chou, 2010; Sauerbrei et al., 2016). Advancing our toolbox such as by developing a comprehensive mutation phenotype map that is community maintainable and in an open format would solve these data requirement problems for genotypic methods. Further, it could play an important role in developing improved antiviral therapies (Chou, 2020; Razonable et al., 2020).

User-friendly tools are also important to enable researchers to make use of such databases (Chevillotte et al., 2010; Hayer et al., 2013). Any such tool built to analyse HHV sequence data should support both sanger consensus sequence outputs, as this is still widely adopted, as well as modern Next Generation Sequencing (NGS) variant outputs (Streck et al., 2023). NGS has become a popular option and provides advantages such sensitivity to low-level variants that helps in early decision making (Chin et al., 2013; Garrigue et al., 2016; Guermouche et al., 2020) and evidence that accumulation of low-level variants may be associated with poor clinical outcome (Houldcroft et al., 2016; Venturini et al., 2022). Additionally there is interest in sequencing wider than just DNA polymerase (HSV1/2:UL30, HCMV:UL54, varicella-zoster virus (VZV):ORF28) and kinase (HSV1/2: UL30, HCMV:UL97, VZV:ORF36) genes as the number of antiviral targets has grown (Kawashima et al., 2017; Lurain and Chou, 2010; Serris et al., 2022). For example HCMV mutants in UL27, UL51, UL56 and UL89 are not sequenced in many diagnostic labs (Kotton et al., 2018).

Here we present an open-source comprehensive database that links HHV mutations to impact on drug sensitivity phenotypes that is interrogatable and maintainable by the community. The database is released along with analytical tools suitable for resistance annotation of modern sequence data from HCMV, HSV1, HSV2, VZV and HHV6.To the best of our knowledge published comparable resources are comparatively affected by issues such as not covering all genes affecting current clinically relevant drugs, inability to update data after publication and closed-source databases that cannot be community verified (Chevillotte et al., 2010; López-Aladid et al., 2019; Lurain and Chou, 2010; Sauerbrei et al., 2016).

### Database Construction

Literature regarding mutation resistance phenotype were identified by a comprehensive PubMed (“Database resources of the National Center for Biotechnology Information,” 2018) search with key terms “Cytomegalovirus [Title] AND resistance [Title/Abstract]”, likewise for the other HHVs. Papers were inspected with regular expressions to detect the occurrence of mutations, from where literature article data was manually extracted. Each entry in the database represents the relationship between a mutation, study, control species, assay method, metadata, and fold change, where fold change values are stored for 11 currently clinically important drugs. Multiple entries may be present for the same variant where either a single publication provides different test methodology to produce independent sensitivity results, or where the same variant is tested across multiple publications. This Fold-change values come in the form of a numeric decimal value if fold changes are possible to extract, or as a string “Resistance” or “Polymorphism” if only a phenotype is described. Study reference data comes in the form of a title, a HTTP link and a DOI. Assay information columns record the method of strain generation, i.e., “marker transfer”, or “isolated strain”, the test method such as “PRA” or “dye uptake assay” and the control strain used to generate a fold change. For some well-studied mutations, co-occurring mutations have been observed to amplify resistance and are recorded where relevant (Chou, 2015; Chou et al., 2016).

Finally, any metadata such as when the entry was created, any notes and a status flag. Only rows with the status “A” for active are returned in the R package and webserver.

It is well understood that a single key mutation is almost always the causative agent that results in drug resistance for the HHV’s, while any other mutations are often neutral polymorphisms. In this database there are entries from isolates with multiple co-occurring mutations, often clinical isolates. We use the following rules when including these entries: a) If the strain is sensitive in vitro, any uncharacterised mutations are accepted as polymorphisms. B) Where a strain has a single uncharacterised mutation, and the others are known polymorphisms, the single unknown mutation is entered with the strains fold change. C) If there are multiple uncharacterised mutations the entry is recorded along with co-mutations and set the status “O” for obsolete, it will not feature in processes. d) If at a later date with new data only a single uncharacterised mutation now is present for an entry as in (c), the entry is re-looked at and treated as in (b).

### Drug resistance genotyping

To facilitate the resistance annotation of virus sequence data we developed Herpes Drug Resistance Genotyping “HerpesDRG”, a R package that enables the assessment of resistance mutations present from variant VCF or FASTA sequence data (Figure 1). FASTA format sequences are aligned to the selected viruses reference genome with MAFFT (Katoh and Standley, 2013) using the “—add –keeplength” parameters, such that both whole genome sequences or genetic fragments (e.g. UL23, UL54) can be inputted. SNP-sites is used to extract SNP’s and downstream functions identify indels (Page et al., 2016). For detection of low frequency mutations Variant Call Format (VCF >= version4.0) and default Varscan2 output file are accepted, where the data has been mapped to the relevant NCBI reference strain (Koboldt et al., 2012). All input formats are converted to a variant which is processed using the VariantAnnotation package to predict their effect on coding regions (Obenchain et al., 2014; R Core Team, 2014). The HerpesDRG “call_resistance” function produces the key output, an annotated variant table including resistance data present in the database. In the case of a mutation present having multiple entries in the database, all relevant entries are returned.

**Figure 1:**
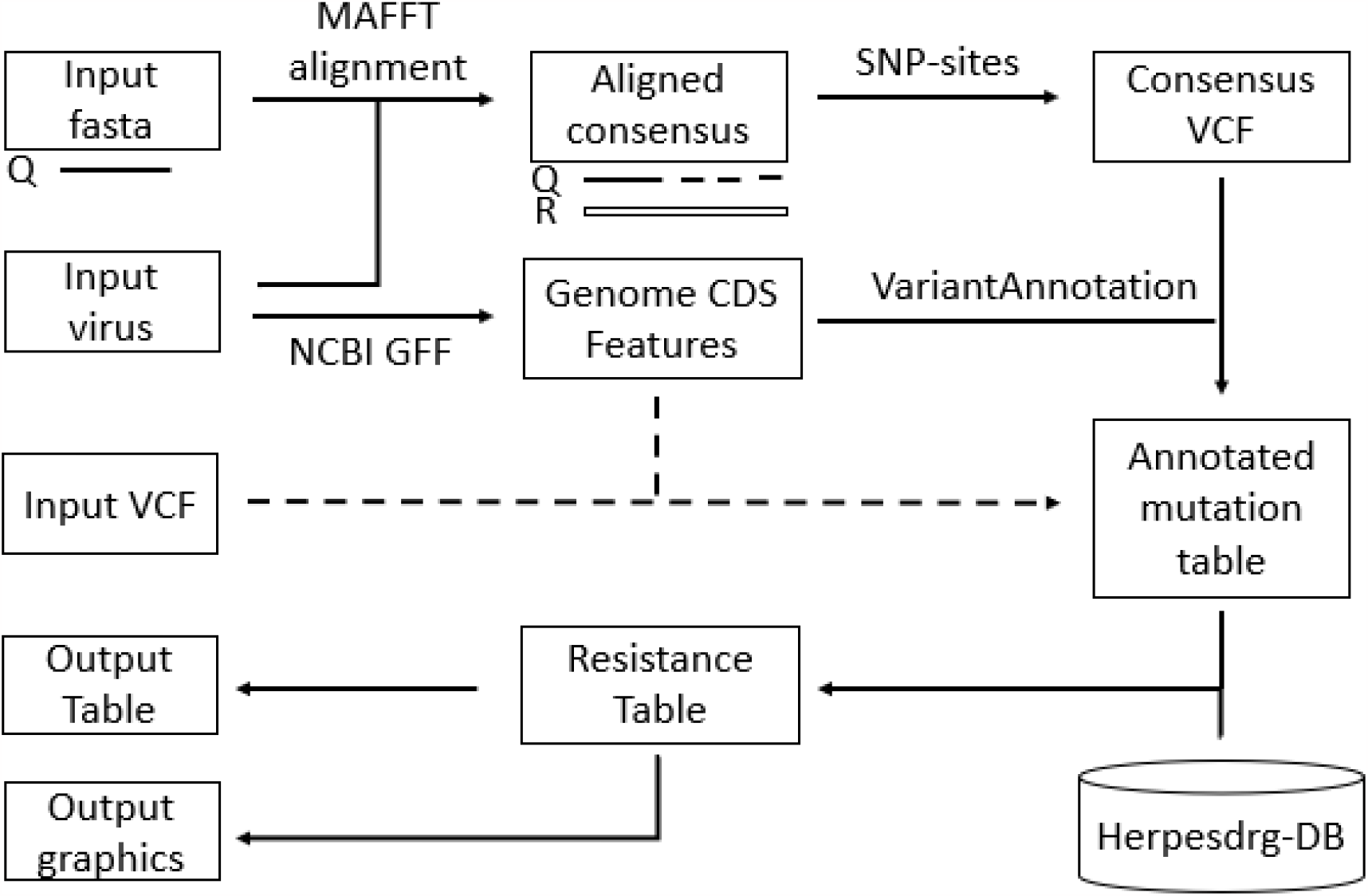
Graphical representation of the HerpesDRG annotation pf FASTA sequence input or VCF inputs (dashed lines). Q – Query sequence, R – NCBI Reference sequence.

A concise clinical output can then be generated from this table using the “make_clin_table” function (Figure 2). Here for each drug HerpesDRG identifies the mutation of maximal fold change found in the sample present at greater than 10% frequency, then allocates a category to that drug accordingly: High level (maximum fold change above 15), Moderate level (maximal fold change between 5-15), Low level (the maximum fold change between 2-5), Polymorphism (less than 2, or recorded as such), Resistant (only anecdotal resistant data was returned), NA (no mutations returned) and evidence strength as: “Good, in vitro”, or NA (no evidence). These fold change cut-off values are in line with recommendations from ‘The Third International Consensus Guidelines on the Management of Cytomegalovirus in Solid-organ Transplantation’ (Kotton et al., 2018) which we use consistently across all viruses.

**Figure 2:**
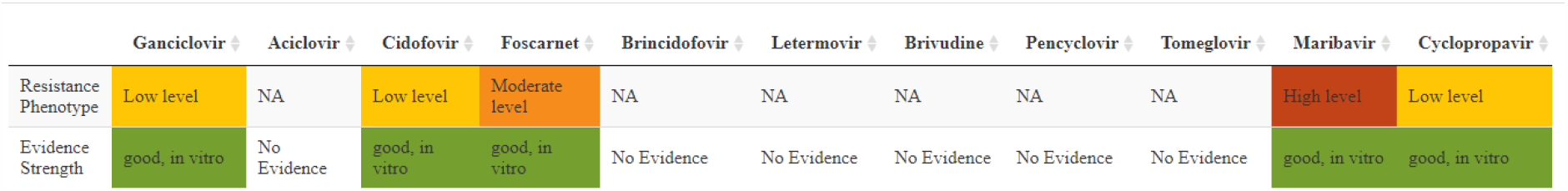
Clinical overview table produced by the “make_clin_table” function for the example data A10.vcf. Example code on GitHub.

### Usage

Install HerpesDRG by running the R command: devtools::install_github(“ojcharles/herpesdrg”). This will take care of installing other dependencies and the database. Inputs for the “call_resistance” function takes as input any of the aforementioned file types, along with which virus to genotype against and flags for whether to return all synonymous / non synonymous mutations (default is only resistance mutants). The above examples were all based on HCMV for consistency but work equivalently for the other herpesviruses.

### User interface

The HerpesDRG toolset is accessible through a user-friendly web interface included in the R package and available at cmv-resistance.ucl.ac.uk/herpesdrg/. Here users upload sequence data as described above and select the virus of interest from the drop-down list. There the full resistance mutations table, the simplified clinical overview and additional graphics are generated automatically. Specific terms of use for the hosted instance are included with the shiny application. The application was developed using the shiny framework (Chang et al., n.d.). We include example files for whole genome analysis (VCF, varscan tab, FASTA), and specific gene regions FASTA.

## Conclusion

We have developed a database that comprehensively captures known drug resistance mutations and their impact on antiviral sensitivity for the antiviral treated Humen Herpes Viruses. The database is released in an open-source format such that it can be community maintained acting as a point of truth for future research and informatic tools. We release this database with an associated tool that allows a low barrier to analysing presence of resistance in sequencing data.

## Funding

O.J.C is supported by the UKRI Medical Research Council grant “MR/N013867/1”. C.V. is funded by Wellcome Trust Grant “204870/Z/16/Z”. J.B. receives funding from the NIHR UCL/UCLH Biomedical Research Centre.

## Conflicts of interest

None declared.

## Acknowledgements

We would like to thank Steven A Kemp and Salvatore Camiolo for their testing of HerpesDRG.

## Notes

http://cmv-resistance.ucl.ac.uk/herpesdrg/

### Competing Interest Statement

The authors have declared no competing interest.

### Summary of Updates

The database has been expanded to address drug resistance mutations in all the antiviral treated herpesviruses, not only HCMV. The manuscript and linked data / code repositories have therefore been updated to reflect this.

